# DReSS: A difference measurement based on reachability between state spaces of Boolean networks

**DOI:** 10.1101/2020.06.19.161224

**Authors:** Ziqiao Yin, Binghui Guo, Shuangge Steven Ma, Yifan Sun, Zhilong Mi, Zhiming Zheng

## Abstract

Researches on dynamical features of biological systems are mostly based on fixed network structure. However, both biological factors and data factors can cause structural perturbations to biological regulatory networks. There are researches focus on the influence of such structural perturbations to the systems’ dynamical features. Reachability is one of the most important dynamical features, which describe whether a state can automatically evolve into another state. However, there is still no method can quantitively describe the reachability differences of two state spaces caused by structural perturbations. DReSS, Difference based on Reachability between State Spaces, is proposed in this research to solve this problem. First, basic properties of DReSS such as non-negativity, symmetry and subadditivity are proved based on the definition. And two more indexes, diagDReSS and iDReSS are proposed based on the definition of DReSS. Second, typical examples like *DReSS* = 0 *or* 1 are shown to explain the meaning of DReSS family, and the differences between DReSS and traditional graph distance are shown based on the calculation steps of DReSS. Finally, differences of DReSS distribution between real biological regulatory network and random networks are compared. Multiple interaction positions in real biological regulatory network show significant different DReSS value with those in random networks while none of them show significant different diagDReSS value, which illustrates that the structural perturbations tend to affect reachability inside and between attractor basins rather than to affect attractor set itself.

**Author summary:** Boolean network is a kind of networks which is widely used to model biological regulatory systems. There are structural perturbations in biological systems based on both biological factors and data-related factors. We propose a measurement called DReSS to describe the difference between state spaces of Boolean networks, which can be used to evaluate the influence of specific structural perturbations of a network to its state space quantitively. We can use DReSS to detect the sensitive interactions in a regulatory network, where structural perturbations can influence its state space significantly. We proved properties of DReSS including non-negativity, symmetry and subadditivity, and gave examples to explain the meaning of some special DReSS values. Finally, we present an example of using DReSS to detect sensitive vertexes in yeast cell cycle regulatory network. DReSS can provide a new perspective on how different interactions affect the state space of a specific regulatory network differently.

## Introduction

Network model is a widely used method to analysis biological systems. There are many kinds of different biological network models are proposed, including protein-protein interaction network, gene co-expression network, gene regulatory network, miRNA network, metabolic network, etc. Researches based on biological network method are mainly focused on two things, structural feature and dynamical feature [31]. The goal of studies which focused on structural features is to explore the key vertexes and society structure of biological systems. This can give us knowledges about the different role of each factor in a system and help us do classification and clustering. The goal of studies which focused on dynamical features is to find the relation between the behavior of biological system and its corresponding outcome. The most representative example for this kind of research is to model the biological regulatory system and predict its outcome. This outcome mostly represents biological phenotype which can be observed.

There is a widely accepted result that cellular regulatory networks are robustly designed [1]. Since then, robustness is one of the most mentioned properties for biological systems [2,3]. Here, the definition of robustness is that the invariance or low variation of a given phenotype when faced with a given incoming variation [4]. Therefore, most calculation methods of robustness are focused on the relation between the perturbation of incoming and the variation of outcoming. Mostly, to biological models based on Boolean networks, the incoming is the initial state. Hence, the perturbation to incoming is the perturbation to the initial state. However, perturbations not only exist in initial state, but in network structure as well. There are both biological and data-related factors can influence the system structure. First, biological factors can affect the biological regulatory interactions. Such as the growth factor can activate the expression of Myc and CyclinD1 [6], and temperature can influence the functional coupling of STIM1 and Orai1 [7] etc. Moreover, there is research pointed out both a passive and active role of DNA methylation to regulatory interactions, where active means changes in methylation can drive changes in regulatory interactions and passive in contrary [8]. Second, networks constructed based on noisy and incomplete biological data are inexact [9]. That is, there may be missing interactions and incorrect interactions in our object networks. There are already methods to assess exploration stage of gene interaction networks [10], which can be used to estimate the possibility of missing and incorrect interactions in our object networks. These researches just focus on the structure itself, but the influence of structural perturbation on systems’ dynamical features are not considered.

Structural perturbations do affect the systems’ dynamical features. To show the affection of structural perturbation to biological system’s stable state, we take fission yeast cell cycle regulatory network as an example. Because fission yeast cell cycle has been studied for years, and there are many widely accepted models that can model the cell cycle process very well [1,12,18]. Here, we use the model from Davidich et al [12] as an example. Boolean network is proposed to model fission yeast cell cycle, which includes 9 vertexes: Start kinase (SK), Cdc2/Cdc13 complex, Ste9, Rum1, Slp1, Cdc2/Cdc13*, Wee1/Mik1, Cdc25 and phosphatase PP. There are two similar vertexes, Cdc2/Cdc13 and Cdc2/Cdc13*. Cdc2/Cdc13 is responsible for the intermediate activity of the complex when Try-15 is in inactive form, while Cdc2/Cdc13* is an indicator of high activity of Cdc2/Cdc13 when Try-15 is unphosphorylated [12]. Each vertex in this network has a state with binary value, 0 or 1. 0 denotes the vertexes is inactive, while 1 denotes the vertexes is active. The states of vertexes are updated in discrete time step under the following function:

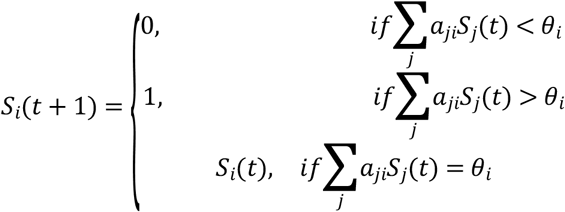

Where *a*_*ij*_ is the elements in network’s adjacency Matrix *A*, *a*_*ji*_ = 1 means there is an activating interaction from vertexes *j* to vertexes *i*, *a*_*ji*_ =− 1 means there is an inhibiting interaction from vertexes *j* to vertexes *i*, and *a*_*ji*_ = 0 means there is no interaction from vertexes *j* to vertexes *i*. *S*_*i*_(*t*) is the state of vertexes *i* at time step *t*. *θ*_*i*_ is the threshold of vertexes *i*, which is set to 0 for all vertexes except Cdc2/Cdc13. Because Cdc2/Cdc13 is positively regulated, so we need to add self-activation to it, which means its *θ*_*i*_ need to be −1. The network is shown in Fig 1.

**Fig 1.**
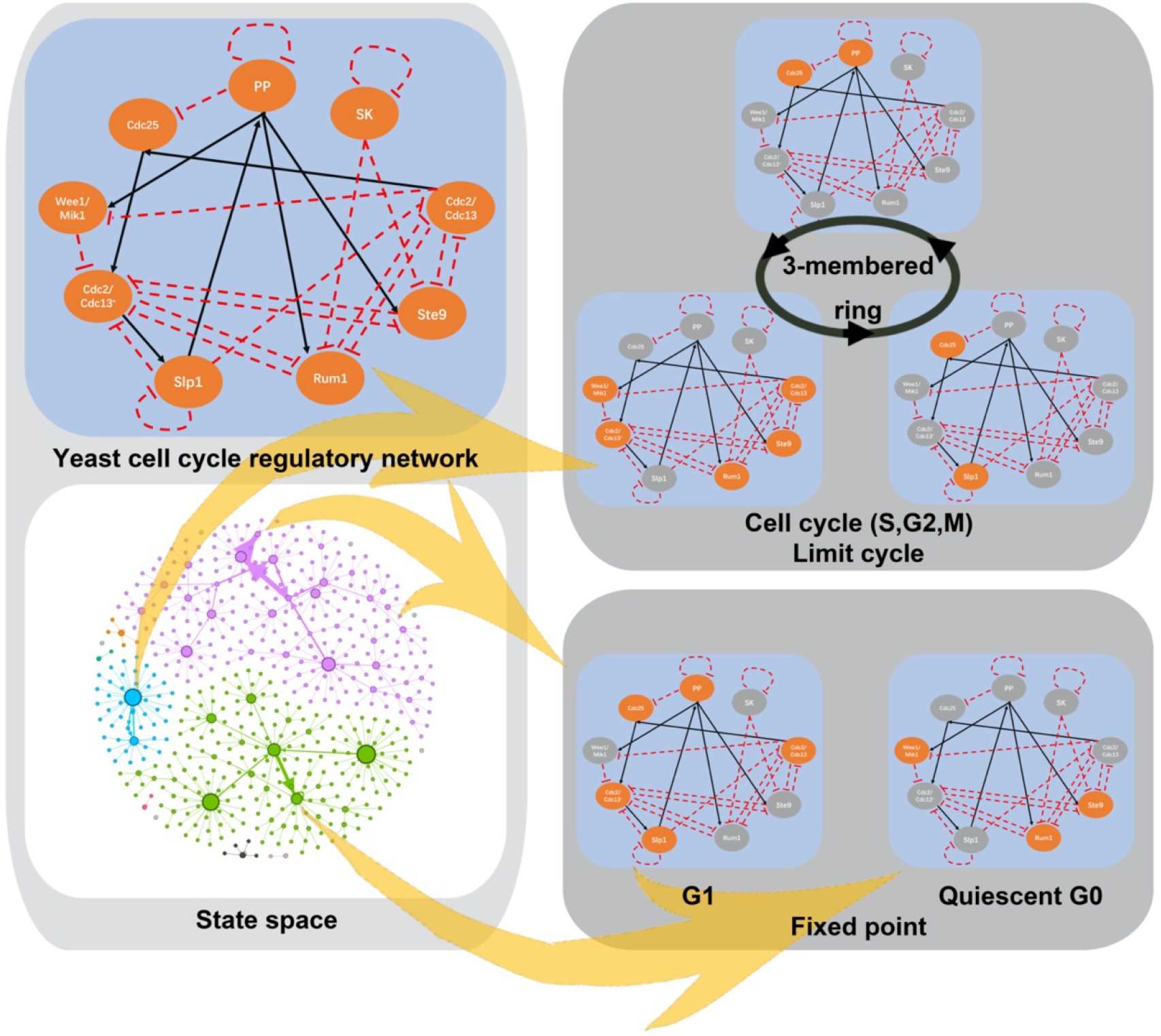
Yeast Cell cycle network. There are 9 vertexes in yeast cell cycle network. Activation interactions are shown in black solid line, and inhibition interactions are shown in red dotted line. Different colors in the state space of yeast cell cycle network represent different connected components, i.e. attractor basins. There are three main attractors in the state space, two fixed points and one limit cycle. Details of these three main attractors are shown in the Fig.

This system has Three main attractors, a limit cycle and two fixed point. Obviously, the limit cycle corresponds to the cell cycle and two fixed point corresponds to G1 state and the quiescent G0 cells. Because there are 9 vertexes with binary value in the network, there are 2^9^ states in the state space. Among them, 490 of 512 states are leads to the three main attractors. The state space of this network is show in Fig 1.

To show structural perturbation can affect network’s state space, three kinds of perturbations are used: delete an exist interaction, add an activation interaction between two vertexes without interaction, add an inhibition interaction between two vertexes without interaction. Without a quantitate index to describe the affection of structural perturbation to the networks state space, we can see its qualitative affection to system attractors. One of the most significant feature of cell cycle system is that there is a special attractor, a limit cycle. However, there are structural perturbations can affect this limit cycle significantly. The influences of structural perturbations to limit cycle are shown in Fig 2. The ones which can change the length of the limit cycle from a 3-membered ring into 2,4,6,8-membered ring are shown in orange, while the ones which can make the limit cycle disappear are shown in green. All boxes in table of Fig 2 are in one of three colors, red, black and grey. Red ones denote to there is a inhibition interaction from the vertex on left to the vertex on top, black ones denote to there is a activation interaction from the vertex on left to the vertex on top, grey ones denote to there is not a known interaction between the vertex on left and the vertex on top. All grey boxes are divided into two part, the upper one denotes to add an activation interaction, and the lower one denotes to add an inhibition interaction. There are 137 kind of one step perturbation to this network. There are 46 of 137 position can make the limit cycle disappear, and 9 of 137 position can change the length of limit cycle.

**Fig 2.**
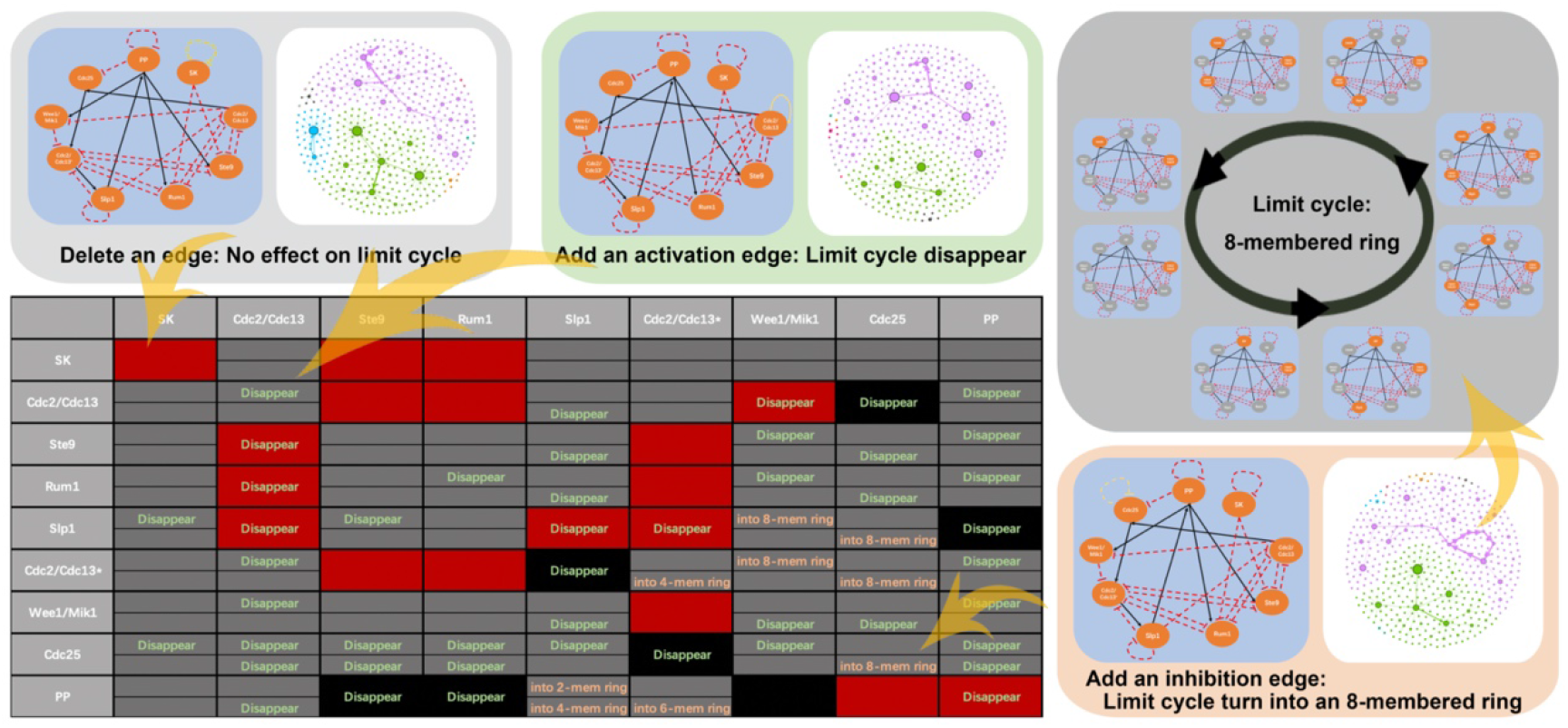
The influence of the structural perturbations of the regulatory network on the limit cycle. There are three colors in the table, red represents there is an inhibition interaction, black represents there is an activation interaction, and grey represents there is no interaction. Perturbations on red and black positions are to remove the original interaction, while perturbations on grey positions are to add a new activation/inhibition interaction. The upper position in a grey box shows the result of adding an activation interaction, while the lower position shows the result of adding an inhibition interaction. The results can be divided into three kinds, limit cycle disappear, change of the length of limit cycle, and no significant change. The results are shown in the box, while the blank positions represent no significant change.

Moreover, these significant changes are not equally distributed to every position. For example, all possible perturbations to interactions from vertex SK cannot make any change to the limit cycle. However, almost every possible perturbation to interaction from vertex Cdc25 can make the limit cycle disappear. That is Cdc25 related interactions are more sensitive to structural perturbation than SK related interactions. Besides, although there are 82 of 137 perturbations that remain the 3-membered ring limit cycle, they still change the state space. Fig 2 gives three examples of each kind of result, disappeaerance of limit cycle, change of the length of limit cycle, and no significant change. The example perturbations are deleting SK-SK self-inhibition, add a CDC2/CDC13 self-activation and add a CDC25 self-inhibition. Compared with the original state space shown in Fig 1, the original limit cycle disappears in two example state spaces. Moreover,

Although there are no significant change in attractor set when SK-SK self-inhibition is deleted, there are still differences between the new state space and the original one. The purple component become larger, while the green component become smaller. That is, part of the states that lead to the purple attractor have changed their destination because of the structural perturbation.

Part of researches which focus on system dynamical features are based on given fixed network structures and do not consider the structural perturbation [11,12]. However, there are researches focused on the influence of perturbations on network structure to the system attractor. As mentioned above, part of the structural perturbation come from the incomplete and noisy data. Therefore, there are researchers trying to recovery the original systems from these incomplete and noisy data. Lü et al proposed a method to predict the position of missing links in a specific network based on the known part [13,14]. Because of the absence of the kinetic parameters, Barabási et al try to predict perturbation patterns from the topology of biological networks [15]. Chao et al tried to evaluate the influence of deleting, adding and switching a specific interaction in a cellular regulatory network [1]. They focused on the influence of structural perturbations to the size change of the largest attract basin, which is an important part of systems’ dynamical feature. Whereas, there is still not a method can evaluate the influence of a specific structure perturbation to the system’s state space quantitively.

Structure perturbation can affect not only the attractor of the system, but the basin of attractor as well. That is, Structure perturbation can change the system’s state space from one to another. A system’s state space is a network, whose vertexes are possible states of the original system. Two vertexes are linked with a directed edge, if one state can transfer into another. In other word, to evaluate the influence of a specific structure perturbation to the system’s state space, is actually to evaluate the difference between two directed networks. There are widely accepted methods to calculate the distance between two networks, such as methods based on maximal common subgraph [16] and based on Laplacian matrix [17]. They consider different feature of networks: similarity, degree distribution, connectivity, etc. However, to state space networks, the most important feature is its reachability, i.e. the ability to get from one state to another within the network. Reachability is one of the most important feature of state space, since it can describe the destination of a specific initial state. Not all stable states are common in biological systems, there are also morbid stable states [32]. Changes of reachability can cause part of the initial states that originally lead to common attractor finally turn into the ones that lead to morbid attractor, which is important in many disease researches. Thus, to evaluate the influence of a specific structure perturbation to the system’s state space, the difference between two directed networks based on their reachability need to be evaluated.

Therefore, DReSS, Difference based on Reachability of State Spaces, is proposed. First, it can quantitively describe the changes of reachability of networks’ state spaces. Second, DReSS obey basic properties including non-negativity, symmetry and subadditivity. Third, a family of index can be defined follow the definition of DReSS to help screen the important interactions (edges) and factors (vertexes) in a system and describe the difference of attractor sets before and after being structural perturbated. In details, diagDReSS is proposed to describe the influence of structural perturbation to systems’ attractor set and iDReSS is proposed to screen the sensitive vertex to structural perturbation. Finally, DReSS can show researchers where they should pay extra attention to in a system to avoid the significant bias on dynamical analysis causing by the potential structural perturbations.

## Result

In this article, basic related concepts are introduced at first. Then, the definition of DReSS is given. Based on the definition of DReSS, two more related index, diagDReSS and iDReSS, are introduced. With such definition, basic properties of DReSS index family including non-negativity, symmetry and subadditivity, etc. are proved in the following part. Next, examples are shown to explain the meaning of DReSS value and its differences against the original network distances. Finally, DReSS index family is applied to real biological network and random networks. Related conclusions are discussed based on the comparation of DReSS distribution on real networks and random networks.

### Definition of DReSS - Difference based on Reachability between State Spaces

Many biological systems are modeled by Boolean networks. A Boolean network *G* = (*V,E*) is a network with Boolean function (details see Methods) as its updating function, where *V* is the set of vertexes and *E* is the set of edges in network. Therefore, each vertex in a Boolean network have a 0-1 state, where 0 represents inhibited and 1 represents activated. In biological systems, vertexes can be DNAs, RNAs, proteins, genes, SNPs, biological factors, etc. And edges are interaction relations, like activation, inhibition, binding, etc, between these vertexes.

The states of vertexes in *G* are continuously updated by updating function until they are stable. At each time step, states of all vertexes form a state array. This state array is the state of *G* at that time step. And all the possible states of *G* and their transfer relations form the state space *S*_*G*_ of Boolean network *G*.

State space *S*_*G*_ is also a network, where all possible states of *G* are its vertexes and the transfer relations are its edges. If *state i* can updated into *state j* in one time step, there is an edge from *state i* to *state j* in *S*_*G*_. State space can describe the transfer relations between states of the original network *G*, which is an important feature of a system. With transfer relations, the final stable states and all intermediate states of any specific initial states can be told. Stable states are always related with stable cellular phenotypes [30,31]. Because only stable states can stable exist and be observed, all unstable states will evolve into stable ones at last. But still, there are lots of perturbations can transfer a stable state into an unstable one, like environmental factors, mutations and external interventions, etc. The unstable state transferred from one stable state not always can return the original stable states but may possibly evolve into another stable states. Therefore, to study the transfer relations between states in state space is very important. However, a state space corresponding one original system. The structural perturbation to the original system would also change the state space as well. Thus, these changes may cause initial states possibly evolve into different stable states before and after the structural perturbation. Here, an index is proposed to describe how much of the initial states changed their destination after a specific structural perturbation. That is, to give an index to describe the reachability differences between the original state space and the new state space.

First of all, A well-defined index to describe the reachability differences between state spaces, especially biological systems’ state spaces, should have its explanation meaning. Second, the index should have basic properties such as commutative, non-negative and obey the triangle inequality etc. Therefore, the definition of DReSS (Difference based on Reachability between State Spaces) is proposed as followed.

To define DReSS, the definition of modified reachability matrix has to be introduced first. Reachability matrix *R* = (*r*_*ij*_)_*N* × *N*_ is a matrix to describe the reachability of networks. *r*_*ij*_ = 1 if there is a directed path from *state i* to *state j* n *S*_*G*_, else *r*_*ij*_ = 0. Particularly, *r*_*ii*_ = 1 for all *i* in *S*_*G*_. A state space *S*_*G*_’s Reachability matrix *R* can be calculated by its adjacency Matrix *A* = (*a*_*ij*_)_*N* × *N*_. *a*_*ij*_ = 1 if there is a directed edge from *state i* to *state j* in *S*_*G*_, else *a*_*ij*_ = 0. And *R* = *I* + *A* + *A*^2^ + … + *A*^*N*^, where *N* is the number of all possible states in state space *S*_*G*_, and operators in the equation are Boolean operators (details see Methods). That is, Reachability matrix *R* is a Boolean matrix, consist of 0 and 1. However, reachability matrix *R* cannot describe the attractor set, which is one of the most important feature of state spaces.

Therefore, the definition of modified reachability matrix 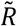 is proposed as followed.

#### Definition 1 Modified reachability matrix

Let *G* = (*V,E*) be a Boolean network with *n* vertexes. *S*_*G*_ is the state space of *G*, and there are *N* = 2^*n*^ possible states in *S*_*G*_. 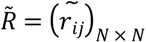 is a *N* × *N* matrix. 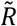 is the modified reachability matrix of *S*_*G*_, if 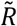 satisfies

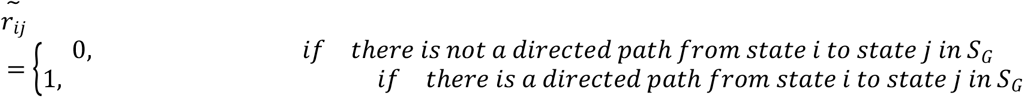

Particularly, 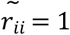 if and only if there is a path from *state i* and can return to *state i* at last including one-step self-loop, otherwise, 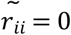.

The only difference between modified reachability matrix 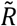 and reachability matrix *R* is the diagonal of the matrix. In the tradition reachability matrix, the diagonal consists of 1 due to *R* = *I* + *A* + *A*^2^ + … + *A*^*N*^. But in state spaces, not all states can reach themselves. Only those states which are attractors or part of attractors can have a path eventually return themselves. Therefore, to state space networks, the original reachability matrix *R* need to be modified into 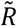, where 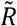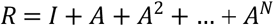. With such a modification, the diagonal of 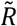 are not all 1 anymore. Only those states in attractors, no matter fixed points or limit cycles, their corresponding position in the diagonal of 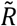 are 1, which can describe the feature of state spaces better than the original reachability matrix *R*. Besides, all states in *S*_*G*_ are *n* dimension Boolean vector. These Boolean vectors, states, can be sorted by its corresponding binary number. That is, there is one and only one unique modified reachability 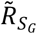 to each state space *S*_*G*_.

With the definition of modified reachability matrix, the Difference based on Reachability between States Spaces can be defined as followed.

#### Definition 2 DReSS

Let *G*_1_ = (*V*,*E*_1_), *G*_2_ = (*V*,*E*_2_) be two Boolean networks with same vertex set *V*. 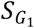 and 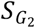 are the state spaces of *G*_1_ and *G*_2_. Let *n* be the number of vertexes in *V*, there are *N* = 2^*n*^ possible states in 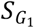 and 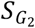. Then, the difference based on Reachability of state spaces (DReSS) between 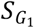 and 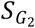 can be defined as

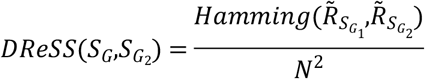

Where 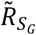 and 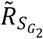 are the modified reachability matrixes of 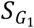 and 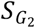, and 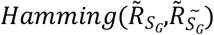 is the Hamming distance of Boolean matrix 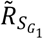 and 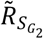.

As *N* = 2^*n*^, *N*^2^ is always a large number in most situations. Mostly, *N*^2^ ≫ *N*. Meanwhile, the number of attractors must less than *N*, and mostly far less than *N*. Therefore, the difference between two attractor sets is a very small fraction of the difference between two whole state space. However, the difference between attractor sets is an important part of the difference of two state spaces. Thus, diagDReSS is introduced to describe the difference between two attractor sets.

#### Definition 3 diagDReSS

Let *G*_1_ = (*V*,*E*_1_), *G*_2_ = (*V*,*E*_2_) be two Boolean networks with same vertex set *V*. 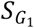 and 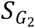 are the state spaces of *G*_1_ and *G*_2_. Let *n* be the number of vertexes in *V*, there are *N* = 2^*n*^ possible states in 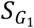 and 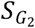. Then, the difference based on Reachability of state spaces (DReSS) between 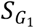 and 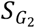 can be defined as

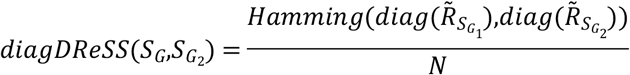

Where 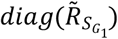 and 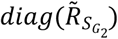 are the diagonal of 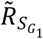 and 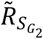.

Although as the definition, DReSS is a difference measurement of state spaces, it can describe the state space difference caused by structural perturbation as well. Therefore, it can be used as an edge centrality at mean time. Because of the heterogeneity of different vertexes and edges in networks, DReSS values are not equally distributed at every possible position of edges. That is, edges with high DReSS values are more likely to be concentrated at specific positions, while the most of other edges are with low DReSS value. Therefore, iDReSS, a vertex centrality based on its surrounding edges’ DReSS value is proposed as followed to describe where structural sensitive perturbations may be clustered at.

#### Definition 4 iDReSS

Let 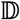 be the set of DReSS of all possible structural perturbation positions in network *G* = (*V,E*). Then, the iDReSS of vertex *i* in *V* can be defined as followed,

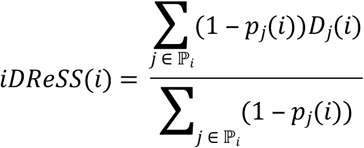

Where ℙ is the position set of all possible perturbations related with vertexes *i*, *D*_*j*_(*i*) is the value of DReSS of perturbation *j*, and *p*_*j*_(*i*) is the p-value of perturbation *j*.

Since iDReSS is a weighted average value of DReSS values of all possible perturbations related with vertexes *i*, iDReSS is also a value between 0 and 1. The average value of *D*_*j*_(*i*) can reflect the structural perturbation sensitivity of vertexes *i* already. But here a weight coefficient (1 − *p*_*j*_(*i*)) is added to it. Perturbations in random networks can also cause affection to the state space. If the significant different structural perturbation sensitive vertexes with biological meanings need to be screened out, those perturbation with significant different DReSS values than those in random networks should be considered at first. Thus, the weight coefficient (1 − *p*_*j*_(*i*)) can help us do this.

Due to the definition of modified reachability matrix 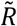,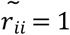 if and only if there is a path from *state i* and can return to *state i* at last including one-step self-loop, otherwise, 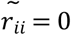. That is, to all states inside the attractor set, their corresponding position in the diagonal of 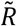 is 1, and all states outside the attractor set, their corresponding position in the diagonal of 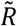 is 0. Therefore, when only the difference between diagonal of 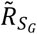 and 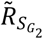 is considered, the index can describe the difference between two attractor sets between 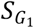 and 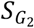. Because the length of the diagonal of 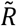 is *N* rather than *N*^2^, so the *N*^2^ is changed into *N* from DReSS to diagDReSS. Besides, as a difference measurement, DReSS obey basic properties below.

#### Property 1 DReSS conform to distance function properties.

To any network *G*, 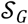 is the set of all possible state space of Boolean networks that have the same number of vertexes with *G*. 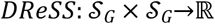, to 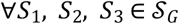, conform to all properties below when network is with deterministic update function and no limit cycle attractor:

a. *DReSS*(*S*_1_, *S*_2_) ≥ 0
b. * *DReSS*(*S*_1_, *S*_2_) = 0⟺*S*_1_ = *S*_2_
c. *DReSS*(*S*_1_, *S*_2_) = *D*(*S*_2_, *S*_1_)
d. *DReSS*(*S*_1_, *S*_3_) ≤ *D*(*S*_1_, *S*_2_) + *D*(*S*_2_, *S*_3_)

And when network is with nondeterministic update function or there is at least one limit cycle attractor, 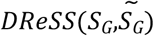 conform to all distance function properties above except (b). Here, deterministic update function means with same input, there will always the same output through the update function, and nondeterministic update function means with same input, there will multiple possible output through the update function.

(a) As 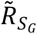 and 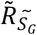 are all *N*^2^ dimension matrix and 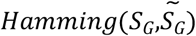 is the total number of difference positions of Boolean matrix 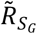 and 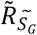, 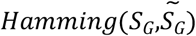 is number between 0 and *N*^2^. That is, 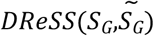 is a number between 0 and 1. Besides, 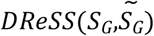 is the percentage of vertexes pairs with different reachability in two networks.

(b) When network is with deterministic update function, there will be only one 1 in the 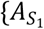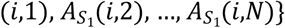. (⟸) if *S*_1_ = *S*_2_, 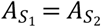. According to the definition, 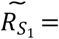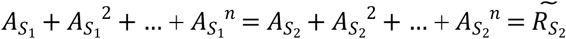. Therefore, 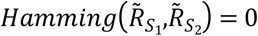. That is, *DReSS*(*S*_1_, *S*_2_) = 0. (⟹) Here, contradiction is introduced to proof *DReSS*(*S*_1_, *S*_2_) = 0⟹*S*_1_ = *S*_2_. Assume that *DReSS*(*S*_1_, *S*_2_) = 0 *and S*_1_ ≠ *S*_2_. As *DReSS*(*S*_1_, *S*_2_) = 0, 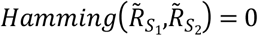. That is, 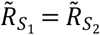. Since *S*_1_ ≠ *S*_2_, there will be at least in elements (*i*,*j*) that 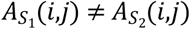. Let’s make 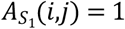, then 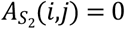. And 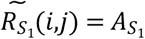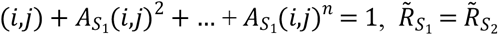, so 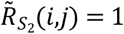. And because 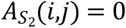, so there is at least one *k*, so that 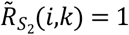 and 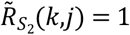. And 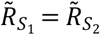, so 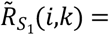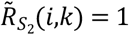 and 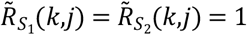. That is, there is a pathway between (*i*,*k*) and a pathway between (*k*,*j*) in *S*_1_. So, state *k* is at downstream of state *i*, and upstream of state *j*. And because 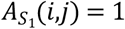, that is state *j* is the next upstate state of state *i*. Therefore, *i*,*j*,*k* formed a limit cycle, or there is more than one upstate state, which contradicts the null hypothesis.

(c) and (d) can be easily proved by the definition of Hamming Distance.

Fig 3 shows two examples that *DReSS*(*S*_1_, *S*_2_) = 0, but *S*_1_ ≠ *S*_2_. Although they are different state space with same 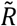, these two state spaces are actually very similar. According to the definition of 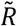, if two state space have same 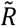, they must have same attractors. If not so, there will be at least one different element on the diagonal of 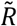. Besides, their reachability must be the same to all intermediate state. These two properties make sure that, from any specific initial state, although the state space may have differences, which makes differences of intermediate states, they always lead to the same attractor (stable state).

**Fig 3.**
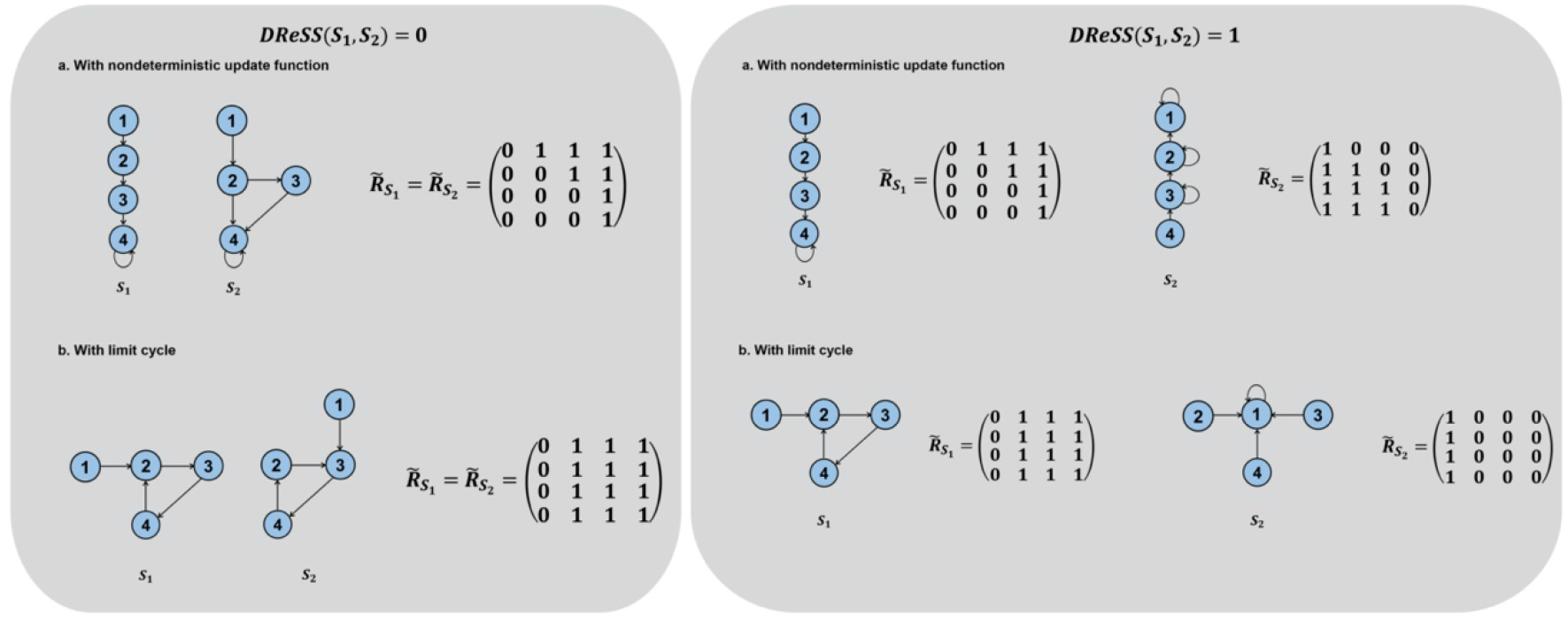
Examples for situations with *DReSS*(*S*_1_, *S*_2_) = 0 and *DReSS*(*S*_1_, *S*_2_) = 1. Each situation is shown with two parts, a. example state spaces for regulatory networks with nondeterministic update function, b. example state spaces for regulatory networks with limit cycle attractor(s).

With such properties, DReSS strengthens its meaning. With *DReSS*(*S*_1_, *S*_2_) = *DReSS*(*S*_2_, *S*_1_), DReSS can ensure there is only one difference value between two specific state spaces, no matter which is the original state space and which is the one after perturbation. With triangle inequality, *DReSS*(*S*_1_, *S*_3_) ≤ *DReSS*(*S*_1_, *S*_2_) + *DReSS*(*S*_2_, *S*_3_), any two state spaces perturbated from the original state space, their differences is smaller than the sum of their differences to the original one.

Although the property of *DReSS*(*S*_1_, *S*_2_) = 0⟺*S*_1_ = *S*_2_ is valid with condition, when it is without the condition, it can still tell a lot of information. First, *S*_1_ = *S*_2_⟹*DReSS*(*S*_1_, *S*_2_) = 0 is always true without any condition. That is, if perturbation cause no change to the state space, the DReSS of the new state space and the original one would be 0. Second, with *DReSS*(*S*_1_, *S*_2_) = 0, *S*_1_, *S*_2_ have same attractor and reachability between states without any condition. Because 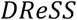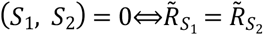, and the diagonal of 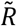 can tell the attractor and the reachability of its corresponding state space. That is, with same 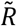 *S*_1_ and *S*_2_ must have same attractor and reachability. Therefore, according to the definition of attractor and reachability, if *DReSS*(*S*_1_, *S*_2_) = 0, from same specific initial state in both state spaces, they can always reach the same attractors.

### Examples when DReSS are exactly 0 and 1

*DReSS*(*S*_1_, *S*_2_) can be exactly 0 and 1, and examples are shown in Fig 3. Obviously, when there is no change in state space, *S*_1_ = *S*_2_, the difference *DReSS*(*S*_1_, *S*_2_) will be exactly 0 according to the definition. However, because the elements in 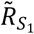 are not independent, that is, 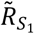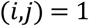 *and* 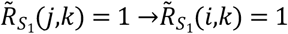, the example for *DReSS*(*S*_1_, *S*_2_) = 1 cannot be constructed without limit. Fig 3 shows an example of *DReSS*(*S*_1_, *S*_2_) = 1. To make the state space ‘totally different’, the path cannot be simply reversed. Modified reachability matrix 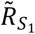 includes two kinds of information, what is the reachability between states and which states are attractors. Therefore, to make *DReSS*(*S*_1_, *S*_2_) = 1, both these two things need to be reversed from *S*_1_ to *S*_2_.

To reverse the reachability between states, there are multiple situations. First, if state *a* is at downstream of state *b* in *S*_1_, state *a* must be at upstream of state *b* in *S*_1_, vice versa. This is the most basic situation. Second, if state *a* cannot reach state *b* and state *b* cannot reach state *a* in *S*_1_, state *a* and state *b* must be in a ring in *S*_2_, vice versa.

To reverse the attractors, the network must with nondeterministic update function when there is more than one attractor in *S*_1_. To make *DReSS*(*S*_1_, *S*_2_) = 1, all states which are not attractors in *S*_1_ must become attractor in *S*_2_, vice versa. When there is more than one attractor in *S*_1_ and the network is with deterministic update function, there is no path between any of two different attractor basins. To reverse this, as mentioned above, any states from one attractor in *S*_1_ must be in a ring with all states in another in *S*_2_. If *S*_2_ is also with deterministic update function, all the states are in a large ring, which against the rule of states in one attractor basin in *S*_1_ cannot reach each other in *S*_2_. And when there is only one attractor, the nondeterministic update function is not a necessary condition. Fig 3 gives an example of *DReSS*(*S*_1_, *S*_2_) = 1 and *S*_1_, *S*_2_ are both with deterministic update function when there are just one attractor in *S*_1_ and *S*_2_.

### Differences between DReSS and traditional network difference measurements

Having proved such properties of DReSS above, the difference between DReSS and traditional network difference measurements should be discussed next. Mostly, graph difference measurements are based on an assumption that all the vertexes in the graph are equal, which makes their order is commutative. However, to biological networks, vertexes are mostly unequal. Different DNAs, RNAs, proteins and other biological complexes, always have their unique function and cannot be replaced by others. And each vertex in state space represents a unique expression pattern of the given biological system. Therefore, two states cannot be considered as the same or similar just because they have same or similar statistical features in state space.

Therefore, different from the former graph difference measurements [16,17], vertexes are considered with order in DReSS. Fig 4 gives an example to show the difference between the former graph difference measurements and DReSS. There are two similar regulatory networks *G*_1_ and *G*_2_. The only difference between them is the position of self-activation interaction. The differences in the original network make their state space different. *S*_1_ and *S*_2_ have same attractors (0,1) and (1,0). In *S*_1_ the rest states are attracted to (1,0), while in *S*_2_ they are attracted to (0,1).

**Fig 4.**
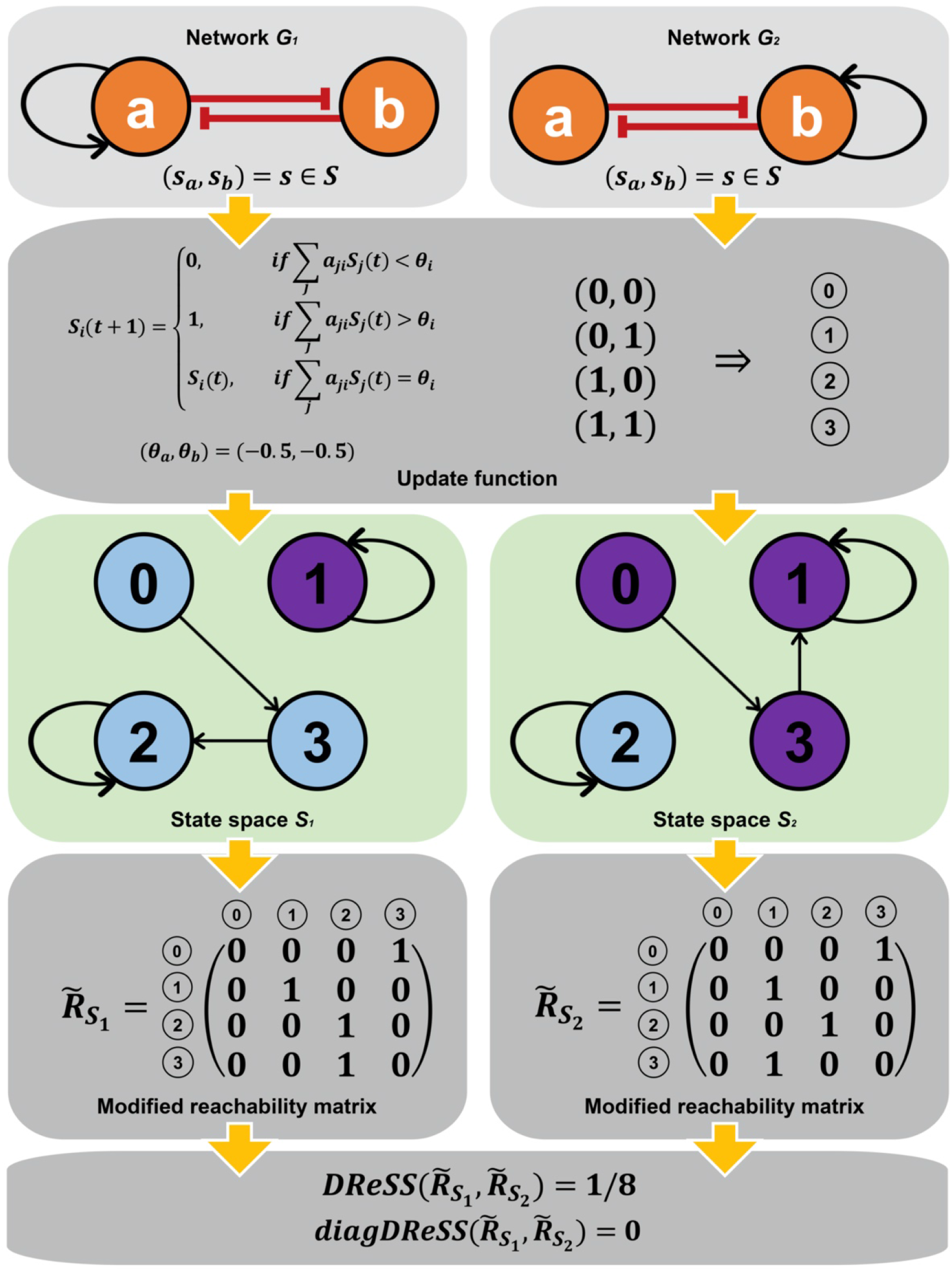
Example for DReSS applied to evaluate the influence of structural perturbation to network’s state space. From the top to bottom, all steps are the original regulatory network, the update function, the state space the modified reachability matrix and the DReSS & diagDReSS value.

There are examples for this kind of mutual inhibition regulatory in biological systems, such as GCR and c-Jun [19] or PTEN and PIP3 [20]. In network science, *G*_1_ and *G*_2_ are exactly the same graph. They have totally the same structure. But vertexes are non-commutative, the conclusion could be totally different. As an example, PTEN and PIP3 have mutual inhibition regulatory relations. PIP3 functions to activate the downstream signaling component including protein kinase AKT, whose isoforms are overexpressed in a variety of human tumors. PTEN, as a phosphatase to dephosphorylate PIP3, always acts the role of tumor suppressor gene. If the mutual inhibition regulatory relations between them is maintained, but give one of them a self-activation, the result would be totally different. Assuming that gene 1 in Fig 4 is PTEN and gene 2 is PIP3, most states in *S*_1_ are attracted to states (1,0), which is the normal states. But in *S*_2_, most states are attracted to states (0,1), which is the cancerous states. Although *S*_1_ and *S*_2_ have the same structural in network science meaning, they lead to totally different results in biological meaning.

However, under the former graph difference measurements, as *S*_1_ and *S*_2_ have totally the same structure, their difference is 0. Although these two state spaces have same attractors and similar state transfer relations, there are still differences between their reachability, that is, 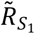 and 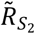 matrix. Thus, their DReSS *DReSS*(*S*_1_, *S*_2_) = 1/8 rather than 0. Because half of the states in state space changed their destination attractor, these two state spaces are not totally the same. This makes the systems more likely to be stable at (0,1) rather than (1,0). Therefore, these two state spaces are same, which shows that DReSS can describe this kind of changes in state space better.

### Data analysis of DReSS in real regulatory networks and random networks

To compare DReSS distribution in the real biological regulatory network and random networks, yeast cell cycle regulatory network is chosen as an example. The structure of real regulatory network is referred from two former related researches [1,12]. The DReSS distribution of all possible structural perturbation to the fission yeast cell cycle regulatory network is calculated. There are three kinds of structural perturbation: delete an existing edge, add an activation edge and add an inhibition edge. To compare the result, the distributions of DReSS under these three kinds of perturbation on three kind of random networks are also calculate: ER random network, BA scale-free network and WS small world network. In random networks, most value of DReSS are clustered around 0, which means structural perturbation has less affection to state space.

However, in yeast cell cycle network, the peak position of DReSS distribution moves toward 1 obviously, and the distribution is more dispersed. There are multiple samples with significant larger DReSS than most samples in random networks. Specifically, there are 2 samples in real regulatory network that with extremely large DReSS value, which are even larger than the maximum value of all 43098 DReSS value among 300 random networks. These two samples are both perturbations toward vertex SK. This result shows that those corresponding structural perturbations make huge difference to the original state space.

On the other hand, there are no sample in real regulatory network with significant smaller DReSS value than the random results except 8 samples with DReSS of exactly 0. All this 8 samples are also structural perturbation towards vertex SK. SK is a special vertex in the original network. It has no input from other vertex, and it only has one self-inhibition as its input. That is, it must be inhibited in any stable state. Thus, it is very sensitive to any input perturbation. SK, Start Kinase, is a simplified vertex as a beginning signal to the system. Thus, it is reasonable that there is rare similar situation in random networks. Meanwhile, there is rare similar situation in real regulatory networks as well. However, this result still reminds us that structural perturbations toward vertex with much more output than input can influence the state space significantly.

Concluded from the results above, DReSS of real biological networks may be more dispersed than the random ones. That is, there are significant sensitive positions and also more stable ones. This result shows that biological network has a clearer role assignment than random network when against structural perturbation. When facing structural perturbations, part of positions in the regulatory network are more sensitive to response, while the rest positions are more stable to keep the system robust, which also shows the importance of DReSS as an index to describe this kind of difference quantitively.

When it comes to diagDReSS, there is no any position with significant different in real biological network compared to random ones. Fig 5.a shows the distribution of diagDReSS of real biological network and random networks. Because there is always only a small part of states can become attractor among of states in state spaces, the diagDReSS distribution is stepped compared to the DReSS distribution. Furthermore, there is no position with significant large diagDReSS value in real biological network compared to random networks, which shows that in both real and random networks, structural perturbation can hardly affect networks’ attractors significantly.

**Fig 5.**
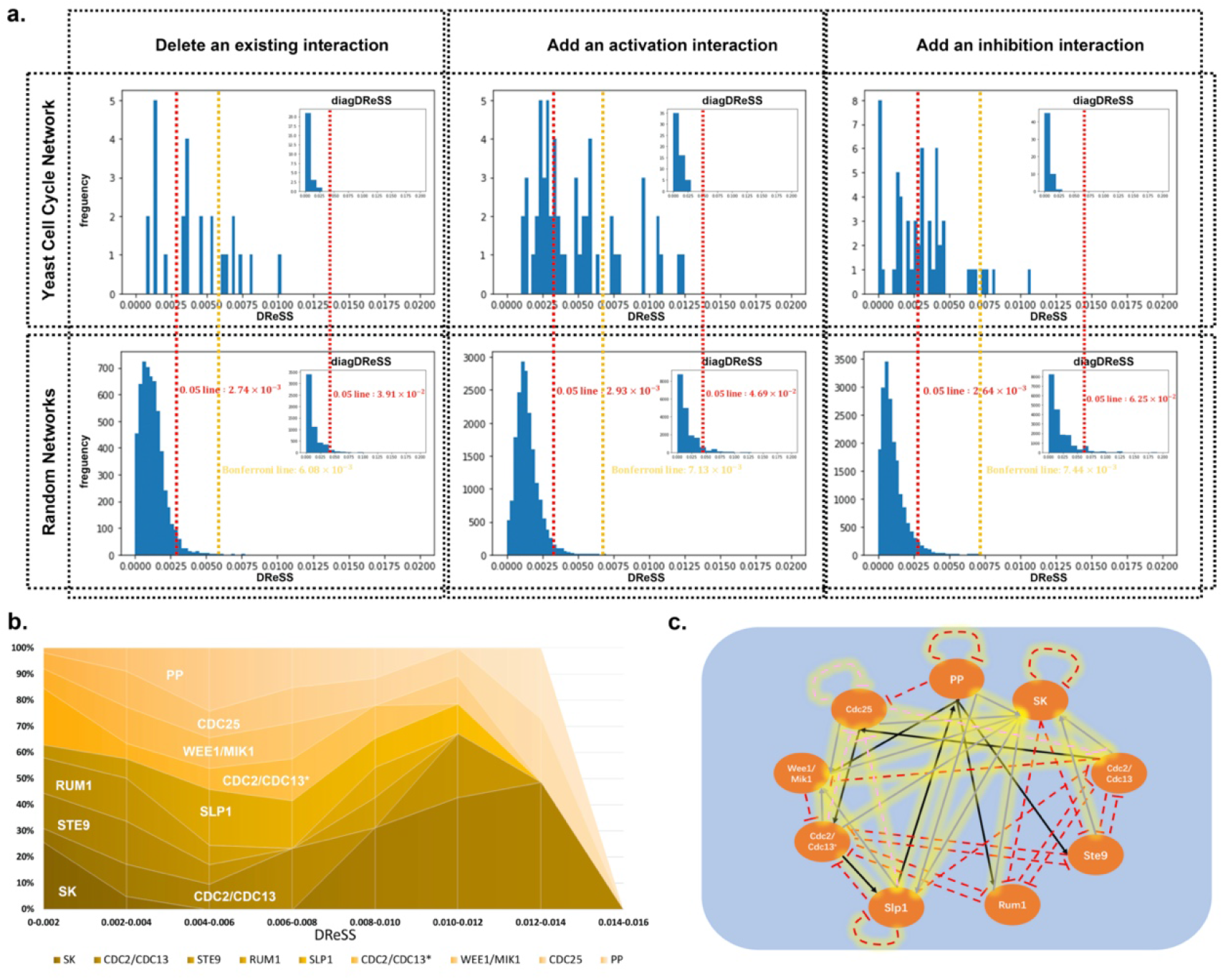
Results analysis on yeast cell cycle regulatory network. a. Distributions of DReSS and diagDReSS in yeast cell cycle network and random networks. The red line is the **5**% significant level threshold, and yellow line is the significant level threshold with Bonferroni correction. b. Distribution of DReSS value of all possible structural perturbation around 9 different vertexes in yeast cell cycle network. c. The structural perturbations with significant difference compared to those in random networks are highlighted in the Fig. Solid lines are activation interactions, while dotted lines are inhibition interactions. The ones with solid color are the perturbations of delete an existed edge, and the ones with translucent are the perturbations of add a new edge.

T-test with Bonferroni correction method (details see Methods) has been introduced to test whether there are samples with significant difference in the yeast cycle network to the ones in random networks. The result shows that there are indeed 22 of 137 possible structural perturbations with significant difference to the ones in random networks, which means perturbation on these positions are more structural sensitive than nearly all the possible ones in random networks. Besides, none of these 137 possible structural perturbations have significant different value compared to the random networks. The positions in real regulatory network with significant different DReSS value from random networks has been light up in Fig 5.c. The structural perturbation positions with significant different DReSS are mostly related with specific vertexes.

Fig 5.b shows DReSS distribution of surrounding possible perturbations of all 9 vertexes in yeast cell cycle regulatory network. The most different vertex is SK whose DReSS clustered around both maximum value and minimum value of the interval. This has already been discussed above. For vertexes like STE9, RUM1 and CDC2/CDC13*, their DReSS is mainly clustered around the lower side of the interval and have no DReSS at top ranks. WEE1/MIK1 and CDC25 are two vertexes whose DReSS are equally distributed in whole interval. That is, when the perturbation is between them and another sensitive vertex, DReSS would be high, and when perturbation is between them and another stable vertex, DReSS would be low. Finally, for CDC2/CDC13, SLP1 and PP, most of their DReSS is clustered at medium and high part of the whole interval, which means perturbations around them may be sensitive. Therefore, iDReSS is calculated for these vertexes in yeast cell cycle regulatory network.

**Table 1.**
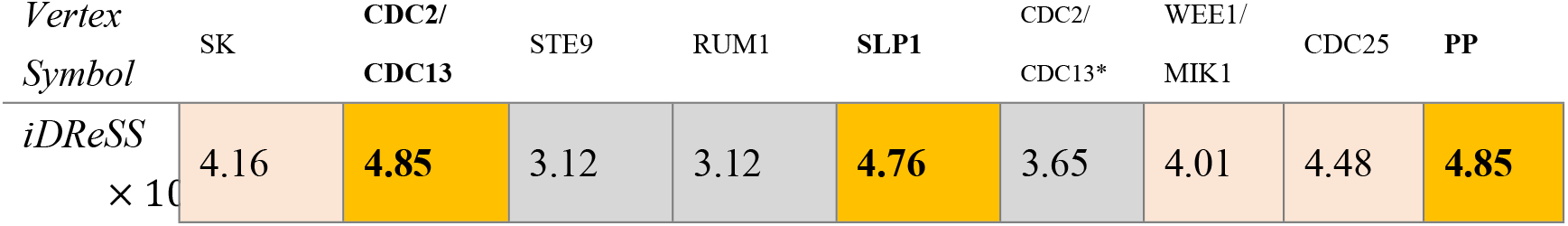
iDReSS value of all 9 vertexes in yeast cell cycle regulatory network.

Table 2 shows the iDReSS value for all vertexes in fission yeast cell cycle regulatory network. Three kinds of vertexes which has been discussed above can be separated under iDReSS. Although SK is the vertex with most high rank DReSS, but it is still not the vertex with largest iDReSS value. This result shows that iDReSS focus on those vertexes whose surrounding positions are mostly sensitive rather than those which have extremely abnormal ones.

All three selected vertexes by iDReSS have their proved biological influence on cell cycle process. CDC genes, Cell Division Cycle genes, are known as key regulatory genes in cell cycle process for years [24]. CDK1 is the protein product of CDC2, whose phosphorylation can lead to the begin of cell cycle. CDC13 can regulates telomere replication, which affects the proliferative capacity of the cell. SLP1 is an effector of the Mad2-dependent spindle checkpoint, which can form a complex with Mad2 essential for the onset of anaphase [25]. PP, phosphatase, can drive protein phosphorylation, which can achieve the widespread reorganization of cellular architecture in mitosis [21].

Moreover, the interactions between these three top vertexes are important as well. CDK1 is the protein product of CDC2. When form a complex with its partner, CDK1 phosphorylation leads to cell cycle progression, which is regulated by PP. Meanwhile, Grallert et al described a PP1–PP2A phosphatase relay which controls the mitotic progression [21]. However, the pathway is blocked in early mitosis due to the inhibition of CDC2 towards PP, and declining CDC2 levels later in mitosis permit PP1 to self-activation again [26,27]. SLP1 is an activator of APC/C (anaphase promoting complex/ cyclosome), which can destroy cyclin B and therefore inhibit Cdk1 (CDC2). These biological results show the importance of interactions between these selected vertexes by iDReSS.

## Discussion

DReSS brings a new quantitively view to difference between state spaces, especially based on structural perturbation. Reachability is one of the most important features for state space, especially for systems with deterministic update function. To systems with deterministic update function, there no path between two attractor basins. That is, the differences of reachability between two state spaces may cause a same initial state to evolve into different stable states in two state spaces. DReSS can show how much of the vertex pairs have changed their reachability between two state spaces. Moreover, besides reachability, attractor set is also an important feature of a state space. diagDReSS can describe the differences of attractor sets between two state spaces. Due to the definition of attractor set of Boolean networks, a state is an attractor of a part of an attractor if and only if it has a path from itself towards itself. Therefore, the differences between the diagonals of two modified reachability matrix 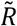 can be used to describe the differences between two attractor sets of two state spaces.Besides, DReSS provides a new method to evaluate the centrality of edges in a graph. There are plenty of researches focus of finding key vertexes of a network. There are many different centrality indexes for vertex, such as degree, betweenness, closeness, eigenvector, k-core, Katz, PageRank, etc. However, there are no so much centrality index for edges compared to for vertexes. Moreover, the edge centralities are mostly based the network structure itself, such as paths, edge betweenness, random walk, information, etc. However, there are rare centrality connecting structural features with dynamical features. DReSS is a measurement that can connect structural features with dynamical features. In this research, only structural perturbation of one edge has been considered. However, DReSS can not only describe the difference between two state spaces caused by only one structural perturbation, it can describe the difference between two state spaces caused by the combination effect of multiple structural perturbation. In biological regulatory networks, changes of environmental factors may cause multiple change to our regulatory system at one time. Meanwhile, the missing and the noisy interactions may not be only one in each system, that is, there may be multiple missing and noisy interactions in a biological system. It is one of DReSS’s advantage that it is with same computational complexity to calculate one or multiple perturbation at one time. Different from the existing edge centralities, DReSS can not only evaluate the centrality of existing edges of a network but can evaluate the potential ones as well. DReSS is actually an index to evaluate the difference between two state spaces. Meanwhile, any structural perturbation can correspond to changes of state space differences. Therefore, DReSS can be used to evaluate the influence of a specific position in a network, no matter there is a known edge at that position or not.

However, the calculation may become a main problem when calculating DReSS of a large system. The computational complexity of 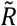 is *O*(*N*^2^) when the system is with deterministic update function, and *O*(*N*^3^) when it is with nondeterministic update function. Meanwhile, here *N* is 2^*n*^ where *n* is the number of vertexes in the original system. Thus, large systems’ 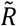 cannot be calculated without simplification such as network module partitions. The average time of calculating a single structural perturbation of yeast cell cycle regulatory network is about 5.82 second and the time of calculating all possible perturbation of yeast cell cycle regulatory network is 797.03 second. When system become larger, the state space will be larger at same time. However, the edges in state spaces are mostly sparse. Furthermore, when the system is with deterministic update function, there will be exactly *N* edges in state space of size *N*^2^ because there is one and only one out-edge for each vertex. That is, there are always much more 0 than 1 in large systems’ 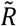. Therefore, these 0 positions may more likely be same in two state spaces, which makes the DReSS of large systems would be much smaller than those of small systems. This can be also explained as when system become larger, the influence of a single structural perturbation to the state space become smaller. Therefore, mostly, the calculation of DReSS value of a single structural perturbation in a large system is not necessary. When the computational complexity exceeds acceptable range, alternative calculating methods such as using random seeds sampling the reachability of state space to get the approximation of DReSS value.

The results above show that there are indeed differences between positions in a network on their sensitivity to structural perturbations. Moreover, these differences are even more significant in real biological networks than those in random networks. This may because the roles of different vertexes in biological systems are clearer and more different than roles of different vertexes in a random network. Based on the results above, part of the interactions are designed to response the changes to the system, while the rest of the interactions are designed to keep the system robust. However, there is not any structural perturbation with significant diagDReSS value in real biological network compared to random networks, which shows the structural perturbations tend to affect reachability inside and between attractor basins rather than to affect attractor set itself. Finally, from the analysis of iDReSS value of real biological network, vertexes can be divided into three groups: the ones aim to keep the robustness of system, the ones aim to make sensitive response to perturbations, and the ones to link the former two group together.

The future research can focus on the theoretical distribution of the DReSS in different situations. With theoretical distribution, the position with significant difference DReSS value can be selected easily. Moreover, the theoretical distribution can help us make a better understanding of the functions of interactions in regulatory systems. Also, the knowledge about the different roles of interactions in a system may help us to find the optimal control method of the system. That is, when DReSS is combined with optimization theory, it can provide new views to find optimal control methods of a system from any initial state evolve into the any stable state with minimum cost. The code can be downloaded at https://github.com/yinziqiao/DReSS-package.

## Methods

In this part, details of basic concepts mentioned in the article are shown to help make a better understanding of the research. The contents are listed in the order in which it appears in the article.

### Boolean function and Boolean network

Boolean function is a kind of function whose arguments assume values from a binary set (usually {0,1}). A Boolean function is a mapping from 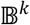 to 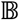, where 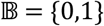. Operation ‘ + ’ mentioned above represents logical operation OR, and ‘ × ’ represents logical operation AND. And the power operations for matrix used above are still based matrix multiplication, whose inside operations are replaced by logical operations. Boolean networks are networks with Boolean function as their update functions. Mostly, the Boolean networks which are used to model biological regulatory networks are directed networks. That is, each edge in the network is with a direction.

### State space

A state space is a set of all possible states of a system. Specifically, to a Boolean network with *k* vertexes, the state space would be 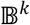, where 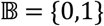. That is, to a Boolean network with *k* vertexes, there are 2^*k*^ states in its corresponding state space. The states can be ordered by their corresponding binary value, i.e. {(0,0,…,0,0),(0,0,…,0,1),…,(1,1,…1,1)}. Besides, as a discrete system, a Boolean network’s state space can be described by a directed graph. States are vertexes in the graph, and there is a directed edge from *s*_1_ to *s*_2_ if and only if to *f*(*s*_1_) = *s*_2_ where *f* is the update function of the system. When the system is with nondeterministic update function, the condition of *f*(*s*_1_) = *s*_2_ turn into T(*s*_1_,*s*_2_) ≠ 0, where T is the transition matrix of the system.

### Attractor and attractor basin

Attractor is a set of numerical values that a system tends to evolve towards. Specifically, to a Boolean network, attractor is a state or a group states, when the system update into such state or such group of states, the system would keep in this state or this group of states unless there are perturbations from outside of the system. The attractor which is a single state called fixed point. And the attractor which is formed with a group of states called limit cycle. The set of all the attractors is the attractor set. Attractors set is the minimal set of states which can only update into states inside the set.

An attractor’s basin of attraction, or attractor basin in short, is set of all the states which will eventually evolve into that attractor. When the system is with deterministic update function, there is no common subset between any two different attractor basins.

### Hamming distance

Hamming distance is a concept from information theory to describe the difference between two string with same length. Its value is the number of positions at which the corresponding symbols are different in two strings. Here, the concept of Hamming distance is borrowed to describe the difference between two matrixes with same size. Its value is the number of positions at which the corresponding elements are different in two matrixes. Thus, its value is between 0 to the size of matrix.

Modified reachability matrix 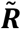

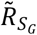 can be simplified into quasi-diagonal matrix 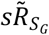 by matrix elementary transformation when network *G* is with deterministic update function, the matrix can be described as:

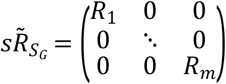

Where *m* is the number of attractors in *S*_*G*_. And when attractor *i* is a fixed point,

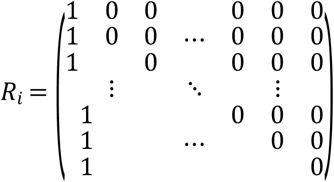

When attractor *i* is a limit cycle,

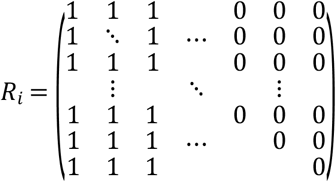

With deterministic update function, one state can have one and only one direct downstream state. Let the basin size of attractor *i* is *L*, then from any state in basin *i*, it takes less than *L* step to reach the attractor *i*. What is more, in a attractor’s basin, there is no ring except the limit cycle. All states in attractor *i*’s basin can reach attractor *i*, and all the states except attractor *i* formed a tree. Therefore, the *R*_*i*_ can be simplified into the form above. And according to the definition, there is no path between attractors. Thus, 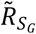 can be simplified into quasi-diagonal matrix 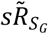 by matrix elementary transformation when network *G* is with deterministic update function. But what need to be noted is that this simplified quasi-diagonal matrix 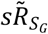 cannot be used for the calculation of DReSS, since the vertexes are ordered in the calculation of DReSS.

### Random networks

In this research,]3 kinds of random networks are used as comparation: ER random network, BA scale-free network and WS small world network. The random networks have to be with same size, same average degree and same activation-inhibition rate with fission yeast cell cycle network. Therefore, random networks are generated of same size and same average degree with yeast cell cycle network by Python package networkx. Then, a probability *p* whose value is the percentage of inhibition interactions in real cell cycle network is set. Finally, edges are turned into inhibition interaction by probability *p*, and keep the rest edges as activation interactions.

### Bonferroni correction

Bonferroni correction is a method to solve the multiple comparison analysis problems. Let *H*_1_, *H*_2_, …, *H*_*m*_ be a family of hypotheses, and *p*_1_, *p*_2_, …, *p*_*m*_ are their corresponding p-value. Bonferroni correction rejects all the null hypotheses if their corresponding *p* ≤ *α*/*m* to control *FWER* ≤ *α*. Specifically, in this research, *m* is the number of all possible position for perturbation in yeast cell cycle networks. The null hypotheses is the DReSS value of perturbation in yeast cell cycle networks obey the same distribution of DReSS value of perturbation in random networks.

